# CACNA2D4 variants are associated with exertional heat stroke susceptibility

**DOI:** 10.64898/2026.01.28.702251

**Authors:** Clara Charpentier, Alioune Fall, Juan De la Rosa Vàzquez, Jonathan Schreiber, Julie Brocard, Patrick Meresse, Dorra Guergour, Caroline Benstaali, Caroline Bosson, Nathalie Roux-Buisson, Bruno Allard, Amy Lee, Isabelle Marty, Julien Fauré

**Affiliations:** Univ. Grenoble Alpes, Inserm, U1216-4, Grenoble Institut Neurosciences, Grenoble, France; Dept of Neuroscience and the Center for Learning and Memory, The University of Texas at Austin, Texas, USA; Univ. Claude Bernard, Institut NeuroMyoGene UCBL – CNRS UMR 5310 – INSERM U1217, Lyon, France; Univ. Grenoble Alpes, CHU Grenoble Alpes, Service de Biochimie, Biologie Moleculaire et Toxicologie de l’environnement, 38000 Grenoble, France; Univ. Grenoble Alpes, Inserm, U1216-4, CHU Grenoble Alpes, Grenoble Institut Neurosciences, Grenoble, France

## Abstract

Exertional Heat Stroke (EHS) is a life-threatening disease defined by severe hyperthermia and sudden neurological dysfunction in healthy individuals exposed to intense physical exercise. As is the case with some forms of malignant hyperthermia, mutations in genes involved in skeletal muscle contraction could predispose some patients to EHS. However, the genetic basis of EHS remains poorly documented. Here we identify EHS-associated variants in the *CACNA2D4* gene encoding the Ca_v_ Ca^2+^ channel subunit, α2δ4, and describe their impact in heterologous expression systems and a knock-in mutant mouse strain heterozygous for the S299R variant (S299R^+/-^). In transfected cells, each *CACNA2D4* variant, including S299R, affected the trafficking and function of α2δ4 and its regulation of the Cav1.1 channel. S299R^+/-^mice showed EHS-like crisis with signs of rhabdomyolysis and an elevation in core body temperature when submitted to an intense exercise protocol. We show that a *CACNA2D4* transcript is expressed in mouse skeletal muscle and that the presence of the S299R variant in α2δ4 induces a modification in the calcium flux triggered by muscle cell depolarization. Altogether, our data reveal an unexpected connection between *CACNA2D4* gene variants and pathophysiological alterations in skeletal muscle Ca^2+^ signaling that may increase EHS susceptibility.

## Introduction

Exertional Heat Stroke (EHS) is a specific and potentially lethal form of hyperthermia occurring during strenuous physical activities in young and otherwise healthy individuals^1,2^. Unlike classical heat stroke, EHS involves skeletal muscles, whose metabolism and contraction provide a source of heat^1,3^. Symptoms include neurologic signs (e.g., confusion, coma, ataxia) and other indicators of physiological distress^1,2^ (e.g., rhabdomyolysis, inflammatory response syndrome and disseminated intravascular coagulation). EHS is a leading cause of death among athletes^4^ and accounts for 36% of all occupational heat-related deaths in construction workers over a 20 years period^5^. EHS incidence was reported as high as 1.2 /1000 in long-distance race runners^6^ and between 0.2 to 10.5 /1000 persons in army forces^7^ . However, the prevalence of EHS is probably underestimated due to difficulties in diagnosis and loss of patient tracking following rapid physical cooling during emergency care^8^.

EHS involves a disequilibrium between thermogenesis and thermolysis during exercise and can be triggered by extreme environmental temperature and humidity^1,9^. However, a specific mechanism explaining the susceptibility of individuals to EHS is still lacking^10^. Like Malignant Hyperthermia (MH)^11^, EHS could be linked to mutations in genes involved in skeletal muscle contraction^12^. MH is triggered by halogenated anaesthetics and manifests as hyperthermia with severe rhabdomyolysis^11^. MH is often caused by mutations in the *RYR1* gene encoding the ryanodine receptor that mediates intracellular Ca^2+^ release and skeletal muscle contraction^13^. However, the prevalence of *RYR1* mutations in EHS is much lower than in MH, suggesting that variants in other genes are involved^14^. Such genes include those encoding proteins involved in Ca^2+^ influx and EC coupling (e.g., TrpV1 and Junctin, respectively) protein involved in EC coupling^15,16^, thus strengthening the hypothesis that both EHS and MH involve dysregulation of skeletal muscle Ca^2+^ signalling^17,18^.

To discover novel risk genes for EHS, we analysed a dataset from exome sequencing of soldiers who underwent a well characterized EHS crisis^15^. In four unrelated individuals, we identified four variants in the *CACNA2D4* gene that encodes the α2δ4 protein which is a subunit of voltage-gated Ca_v_ Ca^2+^ channels^19^. We show that a knock-in mouse strain bearing one of these variants (S299R) variant exhibit EHS crises triggered by an intense exercise protocol. We demonstrated the expression of a muscle specific transcript of *CACNA2D4* and investigated the impact of mutated α2δ4 in the excitation-contraction coupling process.

## Results

### *CACNA2D4* variants in EHS patients

To identify candidate genes involved in EHS, we interrogated an existing dataset from whole exome sequencing of 15 soldiers^15^.These individuals experienced a typical EHS episode with a body temperature ranging from 39.2°C to 41.6°C at the time of the crisis, as well as confusion, disorientation or loss of consciousness^15^. Within this cohort, 14 were diagnosed with MH according to *in vitro* muscle contraction test (IVCT)^13^. We identified three possibly damaging heterozygous variations of the *CACNA2D4* gene (NM_172364.5) in 3 independent patients: c.897C>G; p.Ser299Arg, c.748G>C; p.Gly250Arg and c.486+1G>A. The *CACNA2D4* gene from 26 additional military patients with EHS was further screened by targeted sequencing and the rare c.649G>T; p.Asp217Tyr variant was found in one patient.

All four variants had a low frequency in the genomic databases (Table 1). The c.897C>G; p.Ser299Arg (S299R) and c.748G>C; p.Gly250Arg (G250R) induce substitution of amino acids conserved among all the orthologs of α2δ protein family and were predicted as damaging, whereas the prediction was uncertain for the c.649G>T; p.Asp217Tyr (D217Y). The c.486+1G>A variant was predicted to abolish the consensus splice site which could cause skipping of exon 4. Alternatively, activation of a cryptic site 3-bp upstream could induce the deletion of the last amino acid encoded by the exon (Val162). Notably all variants had potential impacts on the N-terminal part of the α2δ4 protein, and were localized in, or close to, the von Willebrand A (VWA) and Cache domains which are involved in the molecular interaction with the pore-forming α_1_ subunit of Ca_v_ voltage-gated channels^20^ (Fig1A).

**Table 1:** genetic variation in the CACNA2D4 gene (NM_172364.5) found in EHS patients.

| Nucleotide variation | Amino acid variation | MAF | Prediction | Localization | Localization (protein domain) |
| --- | --- | --- | --- | --- | --- |
| c.486+1G>A | - | 1.37e-05 | Inframe deletion of exon4<br>Or p.delV162 * | Intron 4 |  |
| c.649G>T | p.Asp217Tyr | 6.842e-07 | Uncertain /Ambiguous | Exon 5 | VWA_N |
| c.748G>C | p.Gly250Arg | 0.0002 | Damaging/Likely pathogenic | Exon 6 | VWA_N |
| c.897C>G | p.Ser299Arg | 6.158e-06 | Damaging/Likely pathogenic | Exon 8 | MIDAS |

MAF: Minor Allele Frequency (gnomADv4)
Prediction: REVEL/AlphaMissense for missense variants, SpliceAI for intronic variant

For further investigation of their impact, the variants were introduced into the cDNA of human α2δ4 containing an internal HA-tag allowing its detection^21^. When transfected in COS7 or HEK293 cells, the α2δ4 mutant proteins showed normal levels of expression (Fig. 1B). However, compared to the wild-type (WT) α2δ4, the cell-surface levels of the α2δ4 mutant proteins were significantly reduced (Fig. 1B). By immunofluorescence, the S299R variant and other variants were largely retained in intracellular compartments (Sup Fig 1 and data not shown). Along with the auxiliary β subunit, α2δ interacts with and promotes the cell surface trafficking of Ca_v_ channel α1 subunit^22^. Therefore, we next studied the interaction of α2δ4 with the α1 subunit (α1S) of the main Ca_v_ channel expressed in skeletal muscle (Ca_v_1.1). In HEK293 cells, we compared the cell-surface levels of α1S co-transfected with β1a and the α2δ4 variants. Compared to results obtained with the WT α2δ4, the level of α1S that coimmunoprecipitated with the α2δ4 mutant proteins, D217Y and S299R, was significantly lower whereas the other mutations had nominal effects (Fig. 1C). These findings suggest that the *CACNA2D4* mutations have deleterious effects on α2δ4 trafficking to the cell surface as well as its interaction with Ca_v_ channels.

**Fig. 1.**
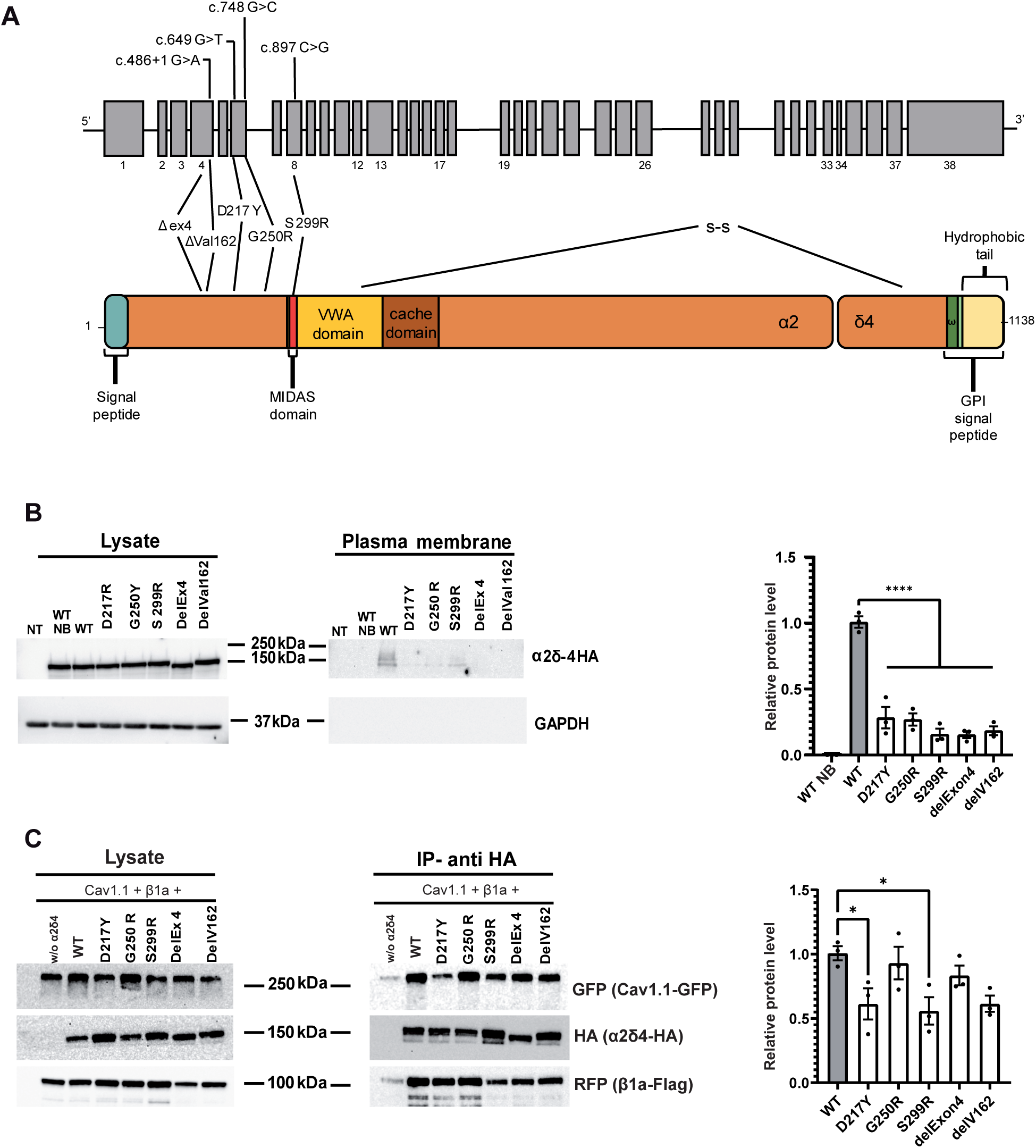
CACNA2D4 variants impair the cell-surface levels of α2δ-4 and its interaction with Cav1.1 A, Schematic of *CACNA2D4* gene and mutations. Upper panel shows exon structure of *CACNA2D4*. Lower panel: domain structure of the α2δ4 protein. Genetic variants and their consequences in the protein are indicated. VWA, von Willebrand factorA; MIDAS, metal ion adhesion site; GPI, glycosyl-phosphatidylinositol anchor site. **B, Impact of *CACNA2D4* variants on cell surface trafficking α2δ-4.** HEK293 cells were transfected with plasmids encoding HA-tagged WT or mutant α2δ-4. After cell surface biotinylation, lysate and plasma membrane fractions were subject to western blotting with anti-HA or anti-GAPDH antibodies. NT= cells not transfected, WT NB= cells transfected with the WT form of α2δ-4 that did not receive biotin during the procedure, WT= WT α2δ-4, EHS variants of *CACNA2D4* are indicated with the consequence on the protein. Graphs shows quantification of the membrane/lysate ratio of HA-α2δ-4 (n=3 ANOVA _✱✱✱✱_,p < 0,001). **C, Impact of *CACNA2D4* variants on interaction with Cav1.1.** HEK293 cells were transfected with plasmids encoding GFP- Cav1.1, RFP-Flag-β1a, and HA-α2δ4 WT or mutant proteins. Lysate and fractions immunoprecipitated by HA antibodies were subject to western blotting with anti GFP, anti-HA, anti-RFP, and anti-GAPDH antibodies. Graph shows quantification of ratio of GFP-Cav1.1 in the immunoprecipitated fraction and the lysate (n=3; _✱_,p=0.0245 for S299R and _✱_,p=0.0495 for D217Y by U-test Mann Whitney).

### *CACNA2D4* expression in skeletal muscle

While initial reports indicated that it is expressed in skeletal muscle^23^, α2δ4 has mainly been studied in the retina where its main function is to regulate the Cav1.4 Ca^2+^ channel in photoreceptors^24–26^. To further analyze the expression of *CACNA2D4* in skeletal muscle, we performed rapid amplification of 5’ cDNA ends (5’RACE) from RNA isolated from mouse quadriceps muscle. Sequencing of the PCR products revealed a *CACNA2D4* transcript composed of all expected exons (NM_001033382.3l), as well as an alternative transcript lacking exons 1, 2 and 16 (Fig. 2A). This second cDNA contained an ORF with an alternate ATG codon in exon 3, producing a α2δ4 protein lacking the first 111 amino acids. Expression of the two transcripts was next tested in mouse skeletal muscle and eye extracts with pairs of primers able to discriminate between both forms. Amplifying exons 14 to 18 revealed 2 products whose size correspond to the presence or absence of exon 16 in skeletal muscle (Fig. 2B). The alternate isoform, which is not expressed in the eye, is referred to hereafter as skm-*CACNA2D4* (encoding skm-α2δ4). When transfected in COS7 or HEK293 cells, the HA-tagged skm-α2δ4 was robustly expressed but was present at the plasma membrane at lower levels than the full length α2δ4 form (Fig. 2C). However, skm-α2δ4 still co-immuoprecipitated with α1S (Sup Fig. 2). Since it lacks exons 1 and 2 and thus the N-terminal signal sequence, skm-α2δ4 may rely on its C-terminal hydrophobic domain to drive its plasma membrane localization. Consistent with this possibility, deletion of the last 35 C-terminal amino acids of skm-α2δ4 (skm-α2δ4ΔCt) caused its accumulation in intracellular compartments (Sup. Fig. 3). These results show that two transcripts of *CACNA2D4* gene are expressed in skeletal muscle, encoding two α2δ4 isoforms that could interact with the α1S subunit of Cav1.1.

**Fig. 2.**
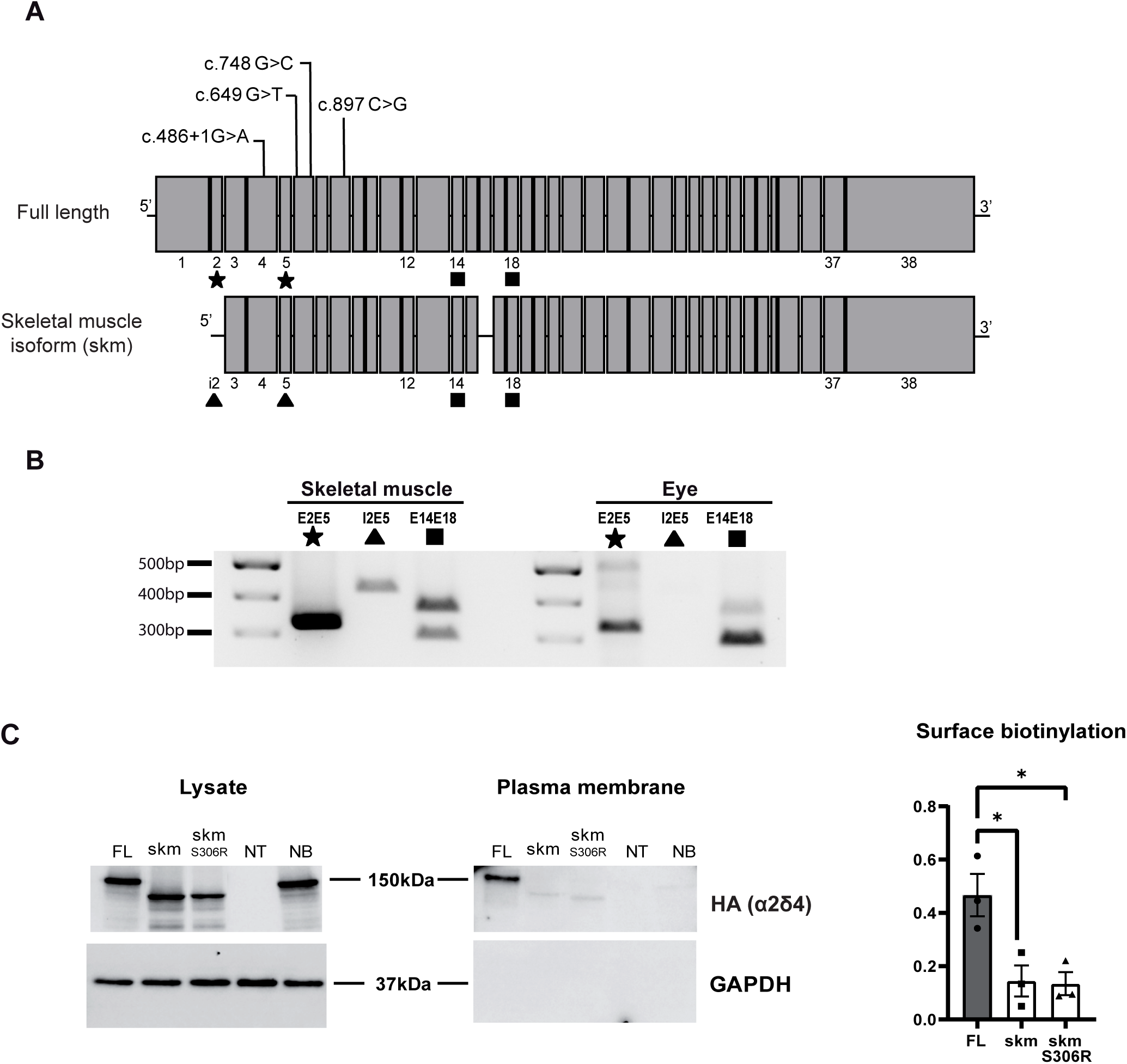
Skeletal muscle expression of the *CACNA2D4* gene. A, Scheme of the Full length (top) and the skeletal muscle isoform (bottom) transcript composition of *CACNA2D4*. Skeletal muscle transcript lacks exon 1, 2, and 16 and was initiated in intron 2 (i2). Numbers refers to the numeration of exons in the gene, i2= intron 2. Symbols indicate the localization of primers used for PCR detection in B. B, Expression of *CACNA2D4* transcripts in skeletal muscle and eye. After RNA extraction and reverse-transcription, cDNAs were submitted to PCR with the indicated pairs of primers. E2E5 primers are localized in exon 2 and 5 symbolized by ★, and is a specific pair for the full length isoform. I2E5 primers are localised in intron 2 and exon 5, and is a specific pair for the muscle isoform symbolized by▴. E14E18 primers are localized in exons 14 and 18 symbolized by ▪, and constitute a pair able to amplify two transcripts with or without a deletion of exon 16. C, Membrane expression of the skeletal muscle isoform transcript of *CACNA2D4*. HA-tagged version of the skeletal muscle or full length isoforms of *CACNA2D4* were expressed in HEK cells. After cell surface biotinylation, lysate and biotinylated plasma membrane fractions were recovered and probed with anti-HA antibody or anti-GAPDH antibody. NT= cells not transfected, NB= cells transfected with the muscle form of α2δ-4 that did not receive Biotin during the procedure, FL= full length form of α2δ-4, skm= skeletal muscle form of α2δ-4, skm S306R = skeletal muscle isoform with the genetic variation S306R. (n=3 ANOVA _✱_,p=0.0211 for skm S306R and _✱_,p=0.0241 for skm).

**Fig. 3.**
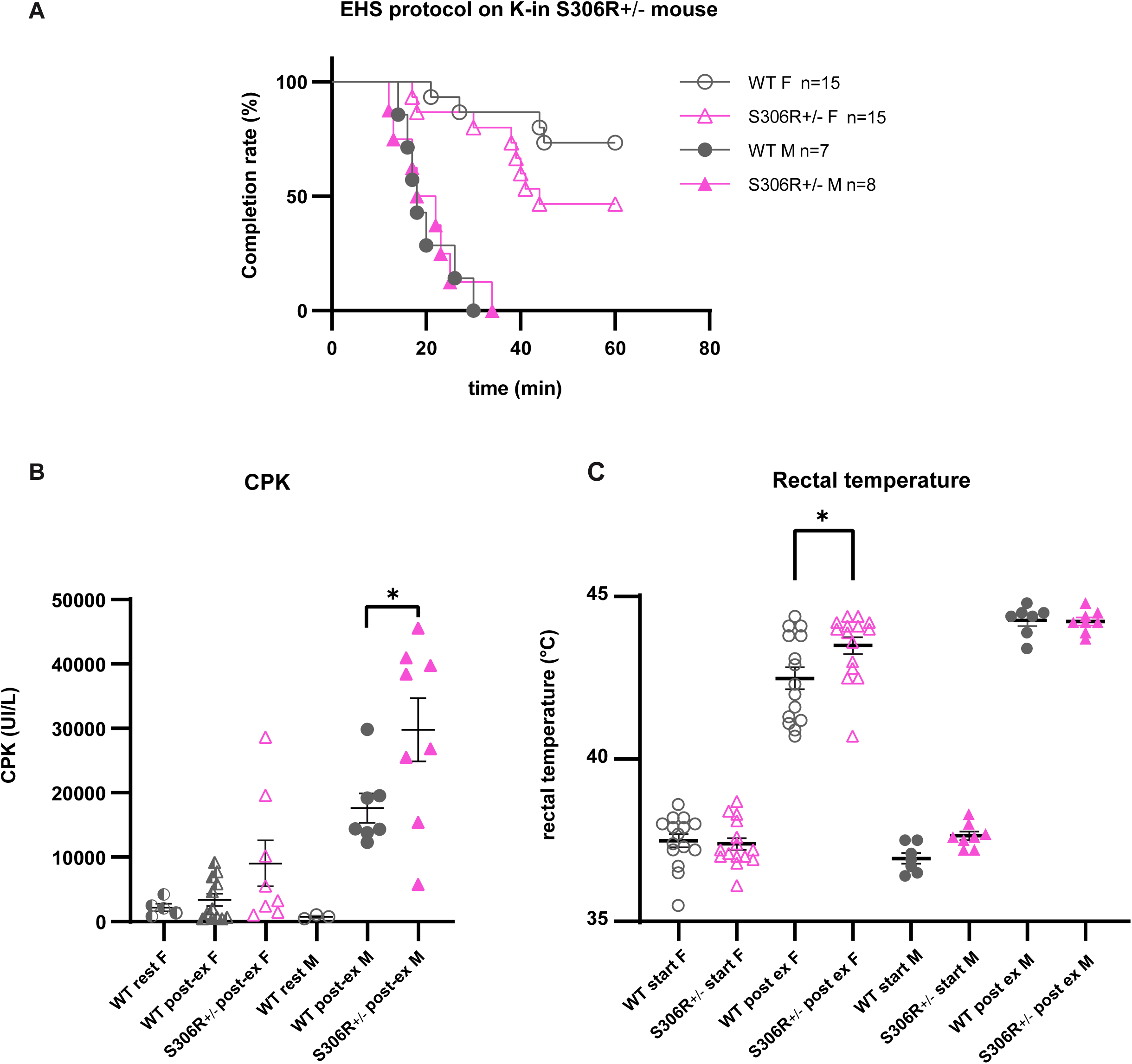
Mouse model with heterozygous S306R mutation reproducing the human S299R variant show EHS-like phenotype. Two months old littermate mice of WT and CACNA2D4-S306R (+/-) genotype were trained 4 days on a treadmill, 10 min per day at the speed of 10 cm/s in environmental conditions. Rectal temperature of the mice was collected daily. After 2 resting days, mice had to sustain a 1h running protocol on a treadmill with speed increasing from 10 cm/s to 20 cm/s with an increase of 5 cm/s per 5 min in an environmental temperature of 39°C and humidity between 20 and 40%. Two endpoints were defined for the exercise: reaching 1h of running or a state described as exhaustion where mice stayed more than 5 seconds immobile without running at the bottom of the treadmill. In any of the 2 situations, rectal temperature was immediately measured, and mice euthanized to collect blood sample. **A,** Percentage of animals reaching any of the 2 endpoints according to the 60min protocol. **B,** CPK plasma level for mice having reached one of the 2 endpoints. Resting CPK levels were collected on animals that have never been submitted to the exercise protocol. Rest= resting CPK level, post-ex = CPK level after mice have reached one of the 2 end points of the protocol. Statistical tests are Unpaired Welch t-test (n > or equal to 8 animals for each genotype and sex) **C,** Rectal temperature was monitored before the exercise (start) or when one of the 2 endpoints was reached (post-ex). Statistical tests are ANOVA (n > or equal to 8 animals for each genotype and sex)

### S306R Knock-in mouse model reproduces EHS phenotype

Serine 299 is a conserved amino acid of the MIDAS domain^27^ of all α2δ subtypes and is involved in the interaction of α2δ with the α1 subunits of Cav channels and the modulation of their electrophysiological properties^28^. To investigate the impact of the EHS-associated S299R mutation, we generated a mouse strain with serine 306 of α2δ4 mutated to an arginine (equivalent to the human S299R variant). The S306R heterozygous (S306R+/-) mice are viable, fertile, and showed no obvious behavioural or physical phenotype during routine care. They were submitted to an intense exercise protocol consisting in a 1 hour run with speed increasing from 10 to 20 cm/s on a treadmill within an environment kept at 39°C and relative humidity between 20 and 40%. Upon reaching the protocol end points (completion of the protocol or exhaustion), rectal temperature was immediately measured, and blood was collected to monitor creatine phosphokinase (CK) levels as a marker of rhabdomyolysis. Male WT or S306R+/- mice were unable to sustain the 1h running exercise (Fig. 3A). They showed high CK levels in their blood (Fig. 3B) and elevated body temperature (Fig. 3C). Interestingly S306R+/- male mice showed a higher level of CK than their WT littermates, indicating more pronounced rhabdomyolysis (Fig. 3B). Most of the WT female mice were able to complete the 1 h run, which was not the case for the S306R+/- female littermates (Fig. 3A). S306R+/- female mice showed a significant higher body temperature (Fig. 3C) at the end of the exercise and a tendency to higher CK levels (Fig. 3B) as compared to the WT littermates. The S306R+/-female mice thus recapitulated cardinal features of an EHS crisis, with exercise-induced exhaustion accompanied by an elevated body temperature and rhabdomyolysis. Of note, S306R+/- male animals showed a massive rhabdomyolysis as compared both to their WT littermates and to the S306R+/- female mice, suggesting the mutation had an impact on their skeletal muscle physiology. These results show that the S306R mutation reproduces an EHS-like phenotype in mice.

### Pathophysiology of *CACNA2D4* S306R variation

To test whether the EHS phenotype of the S306R+/- mice might involve dysregulated Ca^2+^ signalling, we first evaluated the effect of the skm-α2δ4 variants in complex with Cav1.1 channels in transfected HEK293T cells. In these experiments, Ca_v_1.1 was reconstituted with proteins that enable its analysis in heterologous expression systems: α1S, α2δ, the auxiliary Ca_v_ subunit β_1a_, as well as STAC3^29^. We compared the effects of the skm-α2δ4 variants with the full length α2δ4 (FL α2δ4) and the α2δ1 subtype which is highly expressed in skeletal muscle^30^. Consistent with the known modulatory effects of α2δ on Ca_v_ channels^31^, coexpression of α1S with α2δ1 or the full length α2δ4 (FL α2δ4) caused a hyperpolarizing shift in the current-voltage (I-V) relationship of Ca_v_1.1 (Fig. 4A). Compared to channels transfected without α2δ, the voltage of half-maximal activation (V_half_) was ∼10 mV more depolarized for Ca_v_1.1 cotransfected with α2δ1 or FL α2δ4 (Fig. 4B). In contrast, the I-V plots and V_half_ were similar for Ca_v_1.1 transfected without α2δ and Ca_v_1.1 transfected with skm-α2δ4 (Fig.4A and B). Notably, the V_half_ for channels with skm-α2δ4 (37 ± 2 mV) was significantly more depolarized than channels with α2δ1 (25 ± 2 mV). However, the V_half_ for skm-α2δ4 S306R was not significantly different than that for skm-α2δ4 (25 ± 2 mV). Therefore, the skm-α2δ4 variant confers distinct voltage-dependent activation of Ca_v_1.1 compared to α2δ1, but this is not further altered by the S306R mutation.

**Fig. 4.**
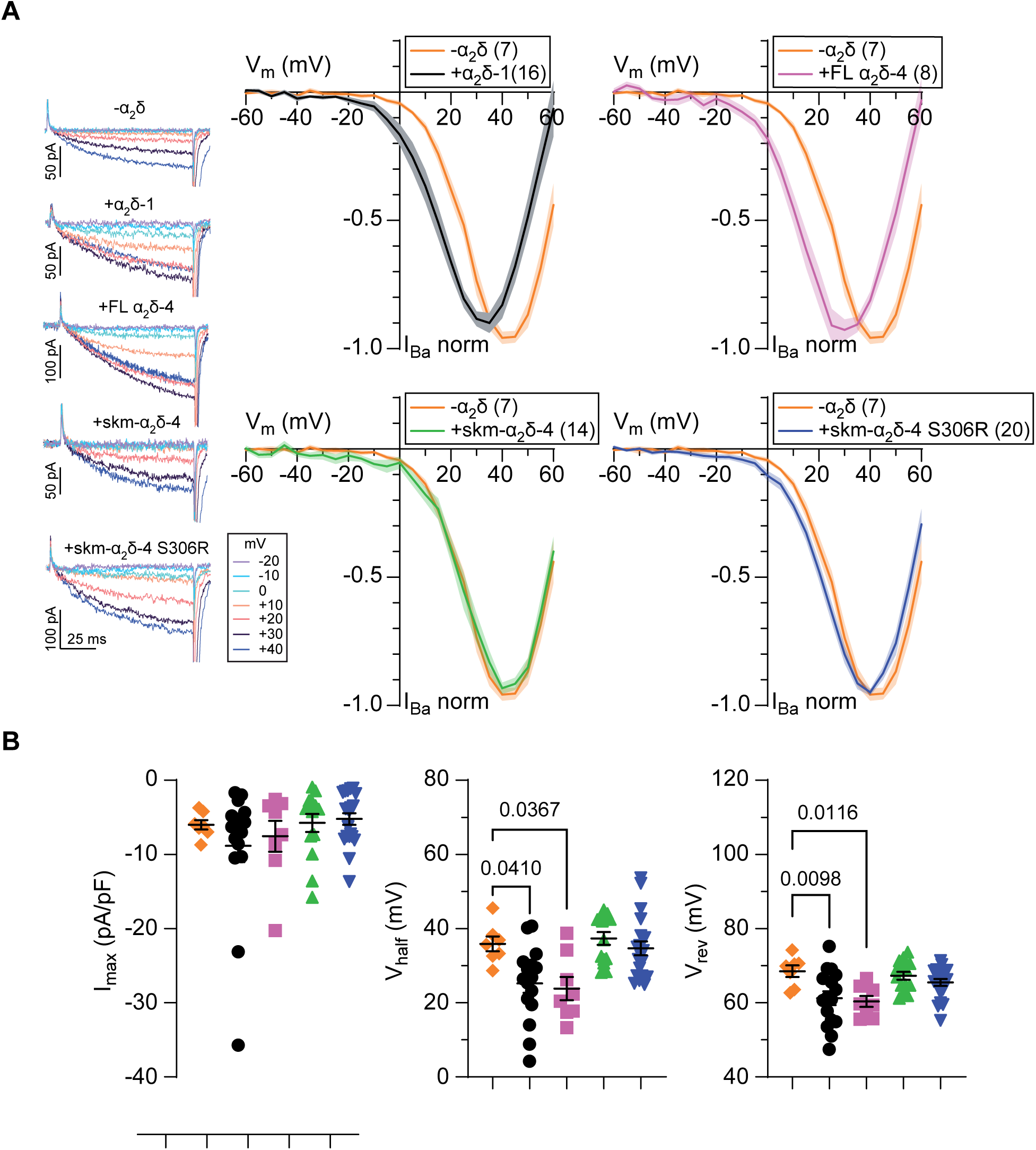
CaV1.1 channels containing skmα2δ-4 exhibit distinct properties compared to channels with α2δ-1. **A**, Representative traces and I-V plots for I_Ba_ elicited by 100-ms pulses from -90 mV to various voltages for CaV1.1 expressed without any α2δ (orange) or with α2δ-1 (black), α2δ-4 (pink), skm-α2δ-4 (green) or skm-α2δ-4 S306R (blue). Data were normalized at the maximum current value on each individual cell and were fit with the Boltzmann equation (smooth line) and plotted with the SEM (shaded region), n values are represented in parentheses. **B**, Maximum current (Imax), voltage of half maximal activation (V_h_) and reversal potential (V_rev_) from Boltzmann fits from the I-V relationships, bars represent mean ± SEM. Kruskal-Wallis test revealed a significant effect of α2δ-1 and FL α2δ-4 on V_half_ (H(4)=20.05, p=0.00051), post-hoc Dunn’s test was used for multiple comparisons. One-way ANOVA revealed a significant effect of α2δ-1 and FL α2δ-4 on V_rev_ (F(4, 60) = 5.101, p = 0.0013), post-hoc Dunnette’s test was used for multiple comparisons.

A physiological effect of S306R could depend on factors expressed in skeletal muscle which are absent in HEK293 cells. To test this, we compared the expression levels and function of Ca_v_1.1 in S306R +/- mice. By Western blot, levels of α1S were higher and Stac3 lower in muscle extracts of S306R+/- compared to WT mice, whereas RYR1 levels were unchanged (Fig. 5A). In electrophysiological recordings of Ca_v_1.1-mediated Ca^2+^ currents, there was trend towards increased current amplitudes in isolated muscle fibres from S306R +/- mice but otherwise the properties of these currents were similar compared to in WT muscle fibers^32,33^. (Fig. 5B). In Ca^2+^ imaging experiments, the SR Ca^2+^ transients of WT and S306R+/- muscle fibers exhibited an early peak followed by a rapid decline toward very low values. Plotting the maximal rates of SR Ca^2+^ release as a function of voltage in each fibre and fitting the relationships obtained with a Boltzmann equation indicated that maximal capacity of SR Ca^2+^ release and voltage of half-activation were not significantly different between genotypes. However, the voltage dependence of the SR Ca^2+^ signals was slightly but significantly steeper in S306R+/- fibers (Fig. 5C). These results show that E-C coupling and proteins involved in this process are perturbed as a consequence of the S306R variant.

**Fig. 5.**
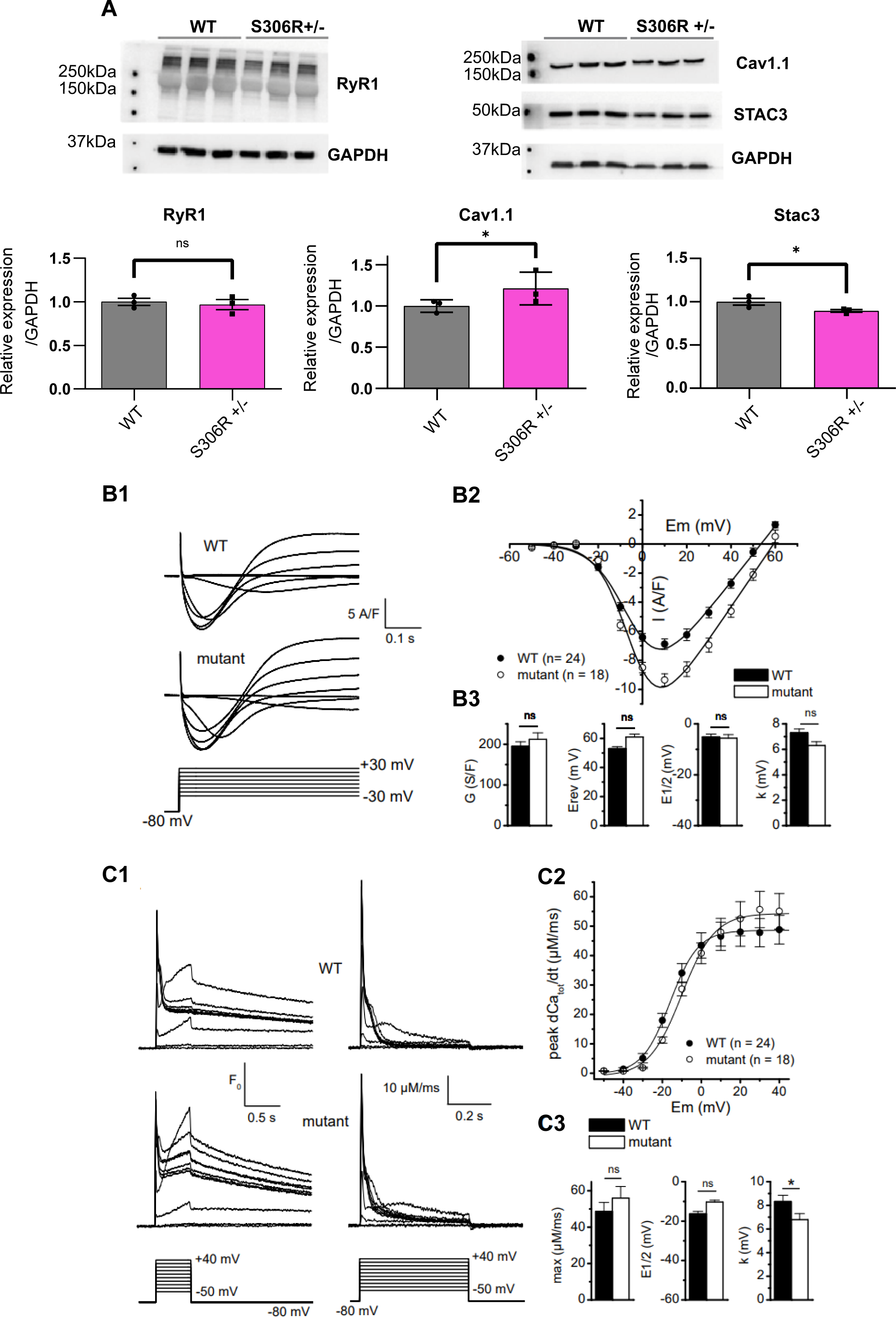
Characterization of muscle fibres from WT and S306R+/- mice A, Expression of RyR1, Cav1.1 and Stac3. Proteins were extracted from Quadriceps muscle from 3 WT or 3 S306R+/- mice and separated by electrophoresis. Each lane represents one extract from an independent animal, antibodies used for detection are indicated as well as molecular weight markers. GAPDH labeling was used as a control and quantification of the level of protein was a ratio of each antibody detection over the GAPDH labeling. N=3 Mann-Whitney U test (*p>0.05) **B, Voltage-gated Ca^2+^ currents in isolated muscle fibres from WT and S306R+/- mice**. B1) Representative Ca^2+^ current traces in the two fibre types in response to the indicated pulse protocol. B2) Relationships between the mean peak values of L-type Ca^2+^ currents and membrane voltage in the two fibre types. B3) Mean of the fitting parameters of current-voltage relationships obtained in each WT and mutant fibre. **C, Voltage-evoked Ca^2+^ transients in isolated muscle fibres from WT and S306R+/- mice**; C1) Representative line-averaged rhod-2 fluorescence changes (left panel) and corresponding rates of SR Ca^2+^ release calculated from the rhod-2 records (right panel) in the two fibre types in response to the pulse protocol shown at the bottom. C2) Relationships between mean values of peak SR Ca^2+^ release flux versus voltage in the two fibre types. C3) Mean (± sem) values for maximum rate of SR Ca^2+^ release (max), voltage of mid-activation (E_1/2_) and slope factor (k) obtained from fitting a Boltzmann function to the peak SR Ca^2+^ release versus voltage relationships in each WT and mutant fibre.

To better resolve the defect in EC coupling in S306R+/- mice, we performed Ca^2+^ imaging on myotubes differentiated from WT or S306R+/- mice. Figure 6A shows that the amplitude of the peak and the total amount of Ca^2^ released upon depolarization was slightly but significantly higher in S306R+/- myotubes as compared to WT myotubes, indicating a modification of EC coupling process in S306R+/- cells. To further assess the role of α2δ4 in EC coupling, we developed a mouse line KO for the *CACNA2D4* gene, inducing a deletion of both full length and skeletal muscle transcripts. The validation of the knock-out was performed by Western blot on proteins extracted from the eye where the endogenous protein can be efficiently detected (Sup. Fig 4). The KO of *CACNA2D4* gene did not alter the viability or the fertility of the mice. Next, we performed Fluo4 calcium imaging upon depolarization in differentiated myoblasts from pups of WT, KO, or S306R+/-, mice. Figure 6B shows that the peak of calcium release, as the area under the curve, in cultured myotubes from *CACNA2D4* KO animals was slightly but consistently reduced, indicating that the absence of the α2δ4 protein induced a perturbation in EC coupling. When myotubes from *CACNA2D4* KO were transduced with lentivirus vectors allowing the expression of the full length or muscle isoforms of α2δ4, the peaks of calcium release were significantly raised, indicating that expression of either form of α2δ4 could rescue the perturbation of EC coupling detected in *CACNA2D4* KO myotubes. Taken together, our findings reveal a novel role for α2δ4 in EC coupling, which is altered in S306R+/- mice.

**Fig. 6.**
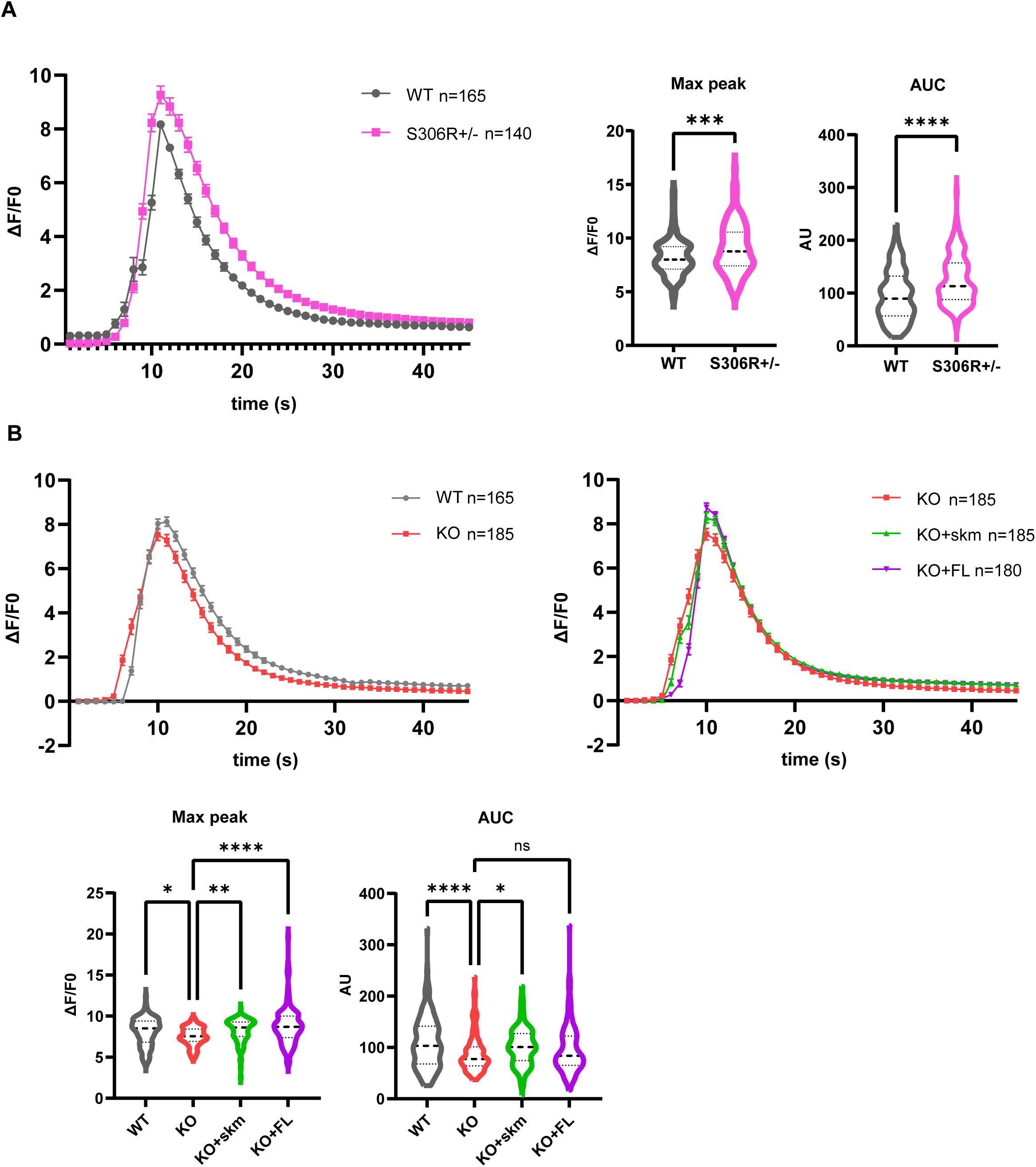
Modifications of EC-coupling in *CACNA2D4*-S306R or *CACNA2D4-KO* mouse myotubes A, Calcium imaging on S306R+/- cultured myotubes. Primary cells were isolated from the heterozygous mouse model *CACNA2D4*-S306R and WT littermates. After 3 days of differentiation, myotubes were stimulated with 140mM KCl in the presence of extracellular calcium using a Fluo4-direct probe. **Left panel: Fluorescence recording during KCl stimulation.** ΔF/F0 represents the ratio of recorded fluorescence over the initial fluorescence for each cell. n= 165 total WT myotubes and n= 140 total S306R myotubes collected on 3 independent experiments. All values are means +/- SEM **Right panel: Quantifications.** Histograms of the means +/- SEM of the peak amplitude for calcium release (Max peak) as well as the area under the curve (AUC). Indicated p values are calculated with t-test on the total number of myotubes N=3, (_✱✱✱_,p=0.0001 for Max peak; _✱✱✱✱_; p<0.0001 for AUC) **B, Calcium imaging on *CACNA2D4* KO cultured myotubes**. Primary cells were isolated from homozygous *CACNA2D4* KO and WT littermates. During the differentiation, cells isolated from KO mice were transduced with lentivirus encoding the full length (KO+FL) or skeletal muscle isoforms of α2δ-4 (KO+skm). After 3 days of differentiation, myotubes were stimulated with 140mM KCl in the presence of extracellular calcium using a Fluo4-direct probe. **Top panels: Fluoresence recording during KCl stimulation.** ΔF/Fo represents the ratio of recorded fluorescence over the initial fluorescence for each cell. n= 165 total WT myotubes and n= 185 total KO myotubes, n= 185 total KO+skm myotubes, and n= 180 total KO+FL α2δ-4 collected on 3 to 4 independent experiments. All values are means +/- SEM. **Bottom panels: Quantifications.** Histograms of the means +/- SEM of the peak amplitude for calcium release (Max peak), area under the curve (AUC) and the slope after the maximum of amplitude (slope). p-values are calculated with an ANOVA on the total number of myotubes. Max peak (_✱✱✱✱_, p=<0.0001; _✱✱_, p=0.0071; _✱_,p=0.0428) AUC (_✱✱✱✱_,p=<0.0001; _✱_,p=0.0101),

## Discussion

In this study, we demonstrate the involvement of *CACNA2D4* missense variants in the EHS phenotype. We identify four missense variants in *CACNA2D4* gene in individuals who experienced well-documented EHS crises. *CACNA2D4* encodes α_2_δ-4, one of four α_2_δ subunits (α_2_δ-1–4) that serve as extracellular subunits of Ca_v_ channels. α_2_δ proteins are glycoproteins produced by post-translational cleavage of a single precursor and linked by disulfide bonds. They regulate Ca_V_ channel trafficking and gating, contributing to excitation–exocytosis coupling, and exert Ca_V_ channel–independent functions^19^. While α_2_δ-1 is the predominant isoform in skeletal muscle^34,35^, the α_2_δ-4 transcript has been identified in skeletal muscle^23^ and was shown to be increased after an EHS episode in mice^36^. While α_2_δ-4 has been primarily studied in the retina^21,25,26,37^, its role in muscle physiology remains poorly characterized. Our findings fit in a genetic landscape where loss of function *CACNA2D4* variants affect the retina and heterozygous point mutations take part in a more complex phenotype, such as the trigger of a hyperthermia crisis during intense exercise in EHS.

### Structural and Genetic Basis of *CACNA2D4* Pathogenicity: From MIDAS Motif Disruption to Aberrant Splicing in EHS

*CACNA2D4* missense variants (p.Ser299Arg, p.Gly250Arg, p.Asp217Tyr) and the splice-site variant (c.486+1G>A) were identified in 4 of 41 unrelated individuals who experienced well-documented EHS crises corresponding to a prevalence of 9.7% in this cohort. This frequency is comparable to the 13% prevalence previously reported for *RyR1* variants^14^ and exceeds that described for other genes associated with EHS or exertional heat illness^16,38^.

The identified variants affect highly conserved residues within functional domains of α_2_δ-4^39,40^. We show that the p.Ser299Arg variant recapitulates an EHS-like phenotype in mice (Figure 3), supporting the pathogenic plausibility of p.Ser299Arg, which targets the MIDAS motif within the VWA domain. Ca^2+^ binding to the MIDAS domain of α2δ is thought to occur while the Ca_v_ channel complex is being assembled in the endoplasmic reticulum, and is necessary for the trafficking of the channel to cell surface^20,27,41,42^. Mutations disrupting the MIDAS sequence impair cation binding, promote intracellular retention, reduce Ca_V_1 interaction, and diminish plasma membrane expression and excitation-exocytosis coupling^27,28,43^. Consistent with these findings, we observed intracellular retention of α_2_δ-4 S306R, decreased surface expression, and reduced interaction with CaV1.1 along with reduced global CaV1.1 levels in transfected HEK293 cells (Figures 1, 2. Sup. fig.1).

Additional evidence for the pathogenicity of the *CACNA2D4* variants derives from disease-associated mutations in homologous residues of other α2δ isoforms. A Gly209Asp substitution in α_2_δ-1, linked to developmental epileptic encephalopathy, causes intracellular retention and loss of Ca_V_ channel trafficking ^40^. Gly209 corresponds to Gly250 in α_2_δ-4, affected by the p.Gly250Arg variant identified in our EHS cohort, and is strictly conserved. Consistent with prior reports, α_2_δ-4 G250R shows reduced surface expression and diminished interaction with Ca_V_1.1 (Figure 1B), underscoring the functional importance of this residue. The c.486+1G>A variant is predicted to disrupt the canonical splice donor site, likely resulting in exon 4 skipping. Although no comparable mutations have been described in other α_2_δ subunits, splice-site alterations account for ∼15% of disease-associated point mutations and commonly lead to aberrant exon skipping^44^. This well-established mechanism further supports a pathogenic link between the c.486+1G>A variant and EHS.

### Context-Dependent Ca^2+^ Dysregulation by *CACNA2D4* variants in EHS

Although α_2_δ-1 is the predominant isoform in skeletal muscle, our findings support a modulatory role for α_2_δ-4 in EC coupling which is altered in S306R^+/-^ mice (Figures 7). Coexpression of skm-α_2_δ-4 causes Ca_V_1.1 to activate at more positive voltages than channels with α_2_δ-1 (Figure 4), as if the channels were no longer modulated by α_2_δ. The S306R mutation would be expected to diminish the contribution of skm-α_2_δ-4 so that a larger fraction of Ca_v_1.1 channels interact with α_2_δ-1, leading to a more hyperpolarized threshold for activation. While we did not observe this in our recordings of Ca_v_1.1 currents in muscle fibers from S306R^+/-^mice (Fig.5), skm-α_2_δ-4 variant may interact with only a subset of Ca_v_1.1 channels in skeletal muscle such that its effect was not possible to resolve. Nevertheless, the trend towards increased Ca^2+^ current amplitudes in the S306R^+/-^ muscle fibers and enhanced E-C coupling in S306R^+/-^ myotubes are both consistent with greater modulation of Ca_v_1.1 by α_2_δ-1 upon loss of function of skm-α_2_δ-4 (Figures 7, 5C). The steeper voltage dependence of SR Ca²⁺ release in S306R^+/-^ myotubes (Figure 6C) suggests a greater cooperativity between voltage sensor activation and RyR1-mediated Ca²⁺ release, perhaps due to the increased dominance of α_2_δ-1 in regulating Ca_v_1.1.

### Sex-Dependent Susceptibility and Extramuscular Roles

Sex-dependent differences observed in our model further support a threshold-based mechanism for EHS susceptibility. Female S306R^+/-^ mice exhibited reduced exercise tolerance and higher body temperatures compared with WT females, whereas males were generally more vulnerable to the protocol, with mutant males showing markedly higher rhabdomyolysis (Figure 3). While controlled experimental studies report minimal sex differences in core temperature responses under standardized workloads and environments, differences in sweating and cardiovascular parameters are evident^45^. In contrast, epidemiological and military data reveal sex-specific patterns of heat illness, with women experiencing higher overall incidence and men showing greater rates of exertional heat stroke^46^. Notably, α_2_δ-4 KO mice display sex-dependent behavioral phenotypes, including impaired motor coordination in females but not males, despite hyperactivity in both sexes^24^, suggesting that α_2_δ-4-dependent circuits or muscle-motor system integration may be differentially buffered. Such sex-specific modulation could influence skeletal muscle or neuromuscular responses to heat and exertion. Collectively, these observations underscore the multifactorial nature of heat illness, in which physiological, behavioral, and environmental factors interact, positioning sex as a context-dependent modifier rather than an independent determinant of thermoregulatory capacity.

Importantly, α_2_δ-4 is expressed beyond skeletal muscle and has been implicated in neuropsychiatric and neurodevelopmental disorders, epileptogenesis, and regulation of locomotor and sensorimotor function^24,47,48^. Subtle alterations in central or peripheral neural circuits controlling locomotion, thermoregulation, or nociception may therefore interact with skeletal muscle Ca²⁺ dysregulation to trigger EHS crises. Collectively, our findings support a model in which the main function of α_2_δ-4 may be to constrain the activity of Ca_V_1.1 channels in skeletal muscle which is compromised in individuals bearing the *CACNA2D4* missense variants. Under conditions of extreme physiological stress, the enhanced function of Ca_V_1.1 and dysregulated E-C coupling could trigger EHS pathophysiology. Our results provide a mechanistic framework that integrates molecular, cellular, and organismal contributors to susceptibility.

## Methods

### Patients Analysis

The cohort of patients and exome analysis was previously described^13^. Genetic analyses were performed after written informed consents were obtained. Whole Exome Sequencing (WES) was performed using Illumina HiSeq© 2000/2500 technology(Illumina, USA). The targeted sequencing of CACNA2D4 gene for 26 patients was performed with a Personal Genome Machine (PGM, Thermofisher Scientific, USA). Libraries were prepared according to the Fast DNA Library Prep Set for Ion Torrent™ manufactuer protocol (New England Biolabs) with a set of specific primers able to amplify human CACNA2D4 exons (NM_172364.5). Individual variation were confirmed by Sanger sequencing.

### Molecular biology

Human CACNA2D4 cDNA (NM_172364.5) with an internal HA tag was already described (Lee et al, 2015). Mouse CACNA2D4 cDNA (NM_001347427.2) was cloned during the RACE PCR, and an internal HA tag was inserted by site directed mutagenesis between amino acids 1512 and 1513. The alternate skm*CACNA2D4* cDNA sequence was submitted to Genbank (accession number : PX837179). Genetic variation found in EHS patients were reproduced in CACNA2D4 cDNA by PCR using specific primers able to reproduce the variation. Cloning procedures were performed with the In-Fusion Snap Assembly Master Mix (638948, TakaraBio). RACE-PCR was performed with the SMARTerRACE5’/3’ Kit (634859, TakaraBio) according to manufacturer protocol. Plasmid and primers used are listed in Table 2

**Table 2.**
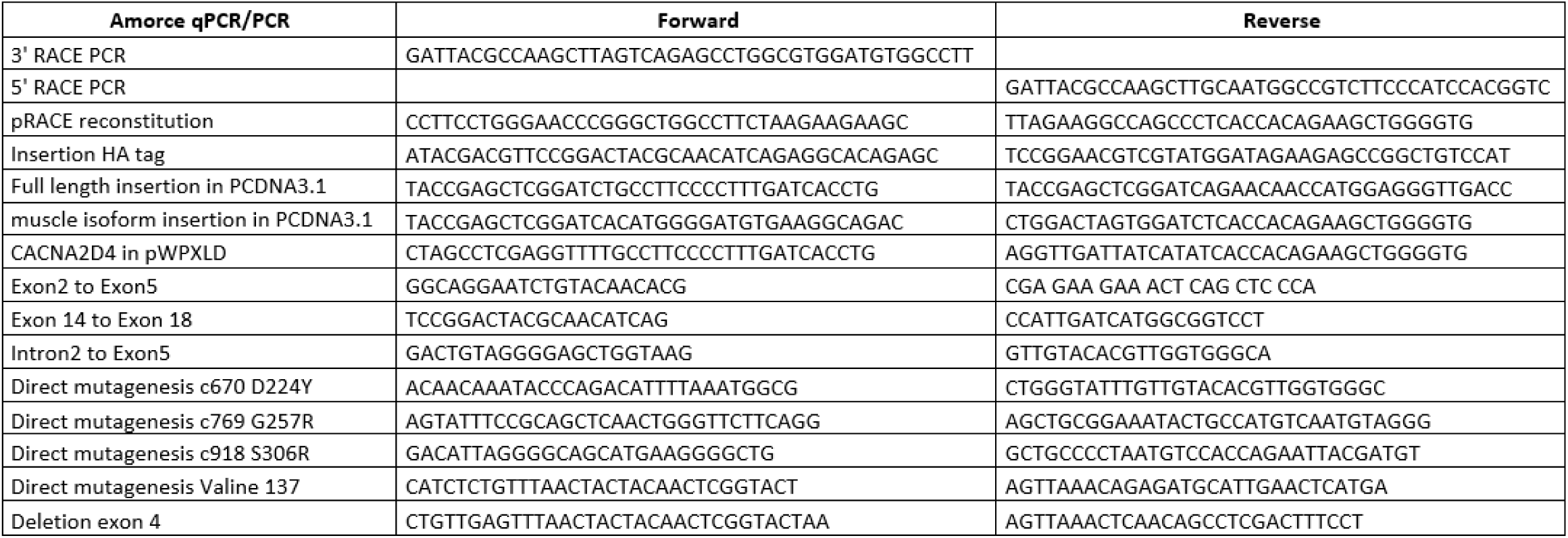
primers.

### Antibodies

Primary antibodies purchased for Western blot or immunofluorescence were as follows: α1s (Sigma HPA048892), Cdk2 (Sant Cruz sc-163), GAPDH (Cell Signaling 2118), GFP (Invitrogen A11122), HA tag (Thermofisher 28183). The antibody against α2δ4 was already described^27^. Secondary antibodies were anti-mouse or anti rabbit antibodies conjugated with HRP (Jakson) or with Cy3 and Alexa488 (Thermofisher).

### Cell culture and transfection

HEK293 and COS7 cells were cultured in Dulbecco’s modified Eagle medium (Pan Biotech P0403500) with 10% fetal bovine serum (FBS), 4.5g/L glucose and 1mM L-Glutamine at 37°C and 5% CO_2_ and were routinely tested for Mycoplasma infections. The day before transfection, cells were seeded on different supports to obtain 70%-80% confluence 24h later. Cell transfections were performed with CalPhos (631312, TakaraBio) or jetPRIME (Polyplus). Media were changed after 24 h and cells were analyzed 48 h after the transfection.

Murine satellite cells were isolated from neonatal mice at no later than postnatal day 2 (P2) as already described^60^. Briefly, after euthanizing the pups by decapitation, dissection of the lower limbs was performed under a binocular to separate all muscles from bones and cartilage. Two enzymatic digestions were then carried out using dispase and collagenase to dissociate the tissue. After two steps of washing and pelleting cells preparation were filtered then through a 40 µm cell strainer into tissue culture dishes containing dissection medium for a pre-plating stage of 3-4h. Non adherent myoblasts were then collected, rinsed, and plated in proliferative medium for 3 to 4 days before being collected and snap frozen at the myoblast stage

### Electrophysiological recordings of transfected HEK293T cells

Human embryonic kidney (HEK) 293T cells, a subclone of the HEK 293 cell line, were acquired from the American Type Cell Culture Collection (ATCC CRL-11268) and maintained in Dulbecco’s modified Eagle’s medium (DMEM, Life Technologies) with 10% of fetal bovine serum (FBS, Gibco) and 1% penicillin/streptomycin (Gibco), at 37 °C in 5% CO_2_. The cells were grown to 70-80 % confluence and co-transfected with cDNAs encoding the human Ca_V_1.1 tagged with GFP, STAC3, β_1a_ and α_2_δ-1 or α_2_δ-4s cDNAs using FuGENE HD transfection reagent (Promega) according to the manufacturer’s protocol. After transfection cells were kept at 37 °C in 5% CO_2_ for 24 h and were dissociated and plated at low density for single cell electrophysiological recordings. β_1a_ and α_2_δ-1 subunits were kindly provided by Kurt Beam (University of Colorado Anschutz Medical Campus).

Whole-cell patch clamp recordings of transfected cells were performed 36-72 h after transfection at room temperature. Data was acquired with an EPC-10 USB patch clamp amplifier driven by PatchMaster (HEKA Elektronik) and analyzed with Igor Pro software (Wavemetrics). Extracellular recording solutions contained (in millimolar): 140 Tris, 1 MgCl_2_, and 20 BaCl_2_. Intracellular solution consisted of (in millimolar): 140 N-methyl-D-glucamine, 10 HEPES, 2 MgCl_2_, 2 Mg-ATP, and 10 EGTA. The pH of intracellular and extracellular recording solutions was adjusted to 7.3, or to the indicated value with methanesulfonic acid. Electrode resistances were typically 4-6 mΩ, and series resistance compensated up to 70%. Leak subtraction was conducted using a P/4 protocol. All average data are presented as the mean ± S.E.M The data were first analyzed for normality using the Shapiro-Wilk tests. Statistical significance differences were determined by ordinary one-way ANOVA and post hoc Tukey’s multiple comparisons test for parametric data and with Kruskal-Wallis test and post hoc Dunn’s tests for non-parametric data, using Prism software (Graph Pad). Normalized current to voltage (I-V) data were fit to the Boltzmann equation (Eq. 1), where *G_max_* is the maximal conductance, *V_m_* is the test voltage, *V_rev_* represents the apparent reversal potential, *V_half_* is the membrane potential at which 50% of the channels are open, and k is the slope factor.

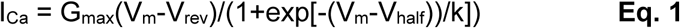

### Calcium imaging

Myoblast from mice were seeded into a drop in the center of a 35 mm dishes coated with Matrigel (1:100 dilution) and maintained in proliferation medium composed of Ham’s F-10 supplemented with 20% FBS, 2% penicillin-streptomycin, and 2% UltroSer G for 3 hours. They were next shifted to differentiation medium composed of DMEM Glutamax-1g/L supplemented with 1% penicillin-streptomycin and 2% Horse serum for 3 days. When required, lentiviral transduction encoding a gene of interest was performed with the induction of differentiation. The medium was then changed the following day, and the cells left to differentiate for 3 more days.To load the cells with the calcium dye, the medium was removed and the cells were incubated with Fluo-4 Direct Calcium assay kit (Invitrogen - Thermo Fischer Scientific #F10471) for 30 min a 37°C. The Fluo-4 was washed with KREBS solution containing 136 mM NaCl, 5 mM KCl, 2 mM CaCl₂, 1 mM MgCl₂, and 10 mM HEPES, adjusted to pH 7.4 and the calcium imaging next performed on a Leica SPE confocal microscope with a 488 nm wavelength of excitation and a 498-600nm PMT collection. The stimulation was performed with a solution containing 100 mM NaCl, 140 mM KCl, 1 mM MgCl₂, 2 mM CaCl₂ in 10 mM HEPES pH 7.4. 40 min time lapse movies were analyse by ImageJ.

### Cell surface biotinylation

HEK293 were seeded at 600 000 cells per 60mm dish, and 24 hours after the transfection, the cell surface biotinylation assay was performed according to manufacturer protocol (A44390, Thermofisher) Kit. Samples of the cell lysates, unbiotinylated and biotinylated fraction were tested next probed by Western-blot, with appropriate antibodies.

### Immunofluoresence

Cos7 cells were seeded at 150 000 cells per 60mm dish containing glass coverslips. 24 hours after transfection, the cells were fixed with a solution containing 4% paraformaldehyde and permeabilized with PBS + 0.1% TritonX100. The blocking solution was made of BSA and Goat serum. Cover Glasses are incubated, with an antibody against HA tag (Monoclonal Antibody Invitrogen) for α2δ4, and a secondary antibody A488.

### Mouse models

The mice models were developed through the Mouse Experimental Biology Platform (PBES) of ENS Lyon, France. The murine models were generated on a C57/Bl6/j genetic background using a CRISPR-Cas9 protocol already described^49^. For the K-in model reproducing the S299R human variation, a single guide RNA guide was designed to target the nuclease cut (sgRNA: GGACATTAGCGGCAGCATGA) and a single-stranded oligodeoxynucleotide (ssODN) used as a template for homology-directed repair to introduce the S306R mutation in the mouse CACNA2D4 gene. Silent mutations introducing a BamHI restriction site for genotyping purposes were introduced in the ssODN and phosphorothioate modifications were added on the 3 nucleotides of 5’ and 3’ ends (ssODN: CGCGATGATGTTCACGAAGTCGTTCTCTCCCAGGGTGTCCAATATGGTGGTAATGGTGT GCTTGGCGATAGCCATCCTCAGCCCCTTCATGGATCCCCTAATGTCCACCAGAATTACGATGTCTTTGGGAGAT). Potential off-target sites for nuclease cut were detected with the CRIPSOR tool. Sites that were mapped less than 20 centimorgans on the same chromosome than the *CACNA2D4* gene were amplified by PCR and sequenced. Founder mice without off-target were selected and backcrossed over WT C56Bl6/j for 4 generations before characterization.

During the selection of founder animals, mice with a single cut due to the sgRNA and no recombination event were analyzed. One animal showed a single nucleotide deletion in the codon of Met341 inducing a frameshift in exon 8 of the *CACNA2D4* gene, generating a *CACNA2D4* KO. Founder animals were backcrossed over 4 generations to create the *CACNA2D4* KO line.

### Exertional heat stroke protocol

This experimental protocol was reviewed and approved by the CEEA-004 ethics committee. All efforts were made to ensure animal welfare in accordance with the 3Rs principle (Replacement, Reduction, and Refinement). The protocol began with a 4-day training period on a BIOSEB treadmill (BX-TM), during which mice ran at a moderate speed of 10 cm/s for 10 minutes per day to acclimate to the treadmill environment. This phase was conducted under ambient temperature and humidity conditions. Simultaneously, rectal temperature was measured at the beginning and end of each session to reduce stress associated with this invasive manipulation. Following a two-day rest period after the training phase, the physical challenge aimed at triggering exertional heatstroke (EHS) was conducted. It consisted of a one-hour running session starting at 5 cm/s, with speed incremented every 5 minutes until reaching 20 cm/s, which was then maintained for the remainder of the session. Environmental parameters were controlled and maintained at 39 °C with humidity between 40% and 60%. Rectal temperature was recorded at the beginning and at the endpoint of the run, once the mouse reached one of the predetermined stopping criteria. Two endpoints were defined: 1-the mouse completes the full one-hour exercise session without stopping, or 2- the mouse remains at the bottom of the treadmill on the grid delivering low-intensity stimuli of 0.5 mA without resuming running for at least 5 seconds, which was defined as exhaustion. Upon reaching either endpoint, the mouse was immediately removed from the treadmill and euthanized. Blood was collected for biochemical analysis. Samples were collected in heparinized tubes and centrifuged to measure serum creatine phosphokinase (CPK) levels.

### Recordings of voltage-gated Ca^2+^ currents and Ca^2+^ transients in isolated mouse muscle fibers

Adult male WT and S306R+/- mouse mice were killed by cervical dislocation before removal of fdb muscles. Single fibers were isolated by a 50 minutes enzymatic treatment at 37° C using a Tyrode solution containing 2 mg/mL collagenase type I (Sigma). Single intact muscle fibers were then released by gentle mechanical trituration of the enzyme-treated muscles in a glass-bottomed experimental chamber, in the presence of culture medium. Prior to trituration, the bottom of the experimental chamber was covered with a thin layer of silicone grease. This enabled single fibers from mouse to be covered with additional silicone so that a 50-100 μm-long portion of the fiber extremity was left out, as previously described. The culture medium solution was replaced by the extracellular solutions (see below). The tip of a glass micropipette filled with an intracellular-like solution containing the Ca^2+^-sensitive dye rhod-2 (see below) was inserted into the silicone-embedded fiber portion. The silver-silver chloride wire inside the micropipette was connected to an RK-400 patch-clamp amplifier (Bio-Logic, Claix, France) used in whole-cell voltage-clamp configuration. Command voltage or current pulse generation was achieved with an analog-digital converter (Digidata 1322A, Axon Instruments, Foster City, CA) controlled by pClamp 9 software (Axon Instruments). The tip of the micropipette was gently crushed against the bottom of the chamber to reduce the series resistance and to allow internal dialysis of the fiber.

Prior to recordings, the rhod-2 dye, diluted at a concentration of 0.1 mM in an intracellular-like solution (see below), was dialyzed into the fiber cytoplasm through the microelectrode inserted through the silicone, within the insulated part of the fiber. Intracellular equilibration of the solution was allowed for a period of 20 min before initiating measurements. Voltage-gated Ca^2+^ currents and voltage-evoked Ca^2+^ transients were recorded simultaneously in voltage-clamped fibers depolarized by 0.5-s-long depolarizing test pulses from −50 to +40 mV. Current changes were acquired at a sampling frequency of 10 kHz. The linear component of the current was removed by subtracting the appropriately scaled value of the steady current measured at the end of a 0.5-s-long −20 mV step applied before each test pulse. The peak Ca^2+^ current, normalized to the capacitance, was fitted in each cell using the following equation: I_Ca_ = G_max_ (E_m_ − E_rev_) / (1 + (exp(E_1/2_ − E_m_)/k)), where I_Ca_ is the peak current density, G_max_ the maximum conductance, E_rev_ the apparent reversal potential, E_1/2_ the half-activation potential and k a steepness factor. Measurements of voltage clamp–activated rhod-2 Ca^2+^ transients were achieved with the line-scan (x,t) mode of a Zeiss LSM 800 confocal microscope with a 50.7-μm-long line scanned every 1.02 ms using excitation from the 561-nm diode laser. Rhod-2 fluorescence transients were expressed as F/F_0_, F_0_ being the pre-pulse baseline fluorescence. Quantification of the Ca^2+^ release flux (dCaTot/dt) from the rhod-2 fluorescence transients was achieved according to previously described procedures^31^. The standard extracellular solution for voltage clamp experiments contained (in mM): 140 TEA-methane-sulfonate, 2.5 CaCl_2_, 2 MgCl_2_, 1 4-aminopyridine, 10 HEPES, and 0.002 tetrodotoxin. The intracellular-like solution contained (in mM): 120 K-glutamate, 5 Na_2_-ATP, 5 Na_2_-phosphocreatine, 5.5 MgCl_2_, 5 glucose, 15 EGTA, 6 CaCl_2_ and 5 HEPES. All solutions were adjusted to pH 7.2.

## Notes

### Competing Interest Statement

The authors have declared no competing interest.

### Summary of Updates

table 2 revised, figures 6 reviserd

### Funding

- Université Grenoble Alpes
- Inserm
- AFM-Telethon

## References

1. Bouchama, A. et al. Classic and exertional heatstroke. Nat Rev Dis Primers 8, 8 (2022).

2. Garcia, C. K., Renteria, L. I., Leite-Santos, G., Leon, L. R. & Laitano, O. Exertional heat stroke: pathophysiology and risk factors. BMJ Med 1, e000239 (2022).

3. Laitano, O., Oki, K. & Leon, L. R. The Role of Skeletal Muscles in Exertional Heat Stroke Pathophysiology. Int J Sports Med 42, 673–681 (2021).

4. Stearns, R. L. et al. Fatal Exertional Heat Stroke Trends in Secondary School Sports From 1982 Through 2022. Sports Health 19417381241298293 (2024) doi:10.1177/19417381241298293.

5. Dong, X. S., West, G. H., Holloway-Beth, A., Wang, X. & Sokas, R. K. Heat-related deaths among construction workers in the United States. Am J Ind Med 62, 1047–1057 (2019).

6. Brillhart, A. et al. Exertional Heat Stroke at the Vermont City Marathon, 2012 to 2023: High Incidence Despite Spring Season in the Northern United States. Clin J Sport Med 10.1097/JSM.0000000000001367 (2025) doi:10.1097/JSM.0000000000001367.

7. Alele, F. O., Malau-Aduli, B. S., Malau-Aduli, A. E. O. & J Crowe, M. Epidemiology of Exertional Heat Illness in the Military: A Systematic Review of Observational Studies. Int J Environ Res Public Health 17, 7037 (2020).

8. Flouris, A. D., Notley, S. R., Stearns, R. L., Casa, D. J. & Kenny, G. P. Recommended water immersion duration for the field treatment of exertional heat stroke when rectal temperature is unavailable. Eur J Appl Physiol 124, 479–490 (2024).

9. O’Connor, F. G. Heat-Related Illnesses. Ann Intern Med 178, ITC97–ITC112 (2025).

10. Roberts, W. O. et al. ACSM Expert Consensus Statement on Exertional Heat Illness: Recognition, Management, and Return to Activity. Curr Sports Med Rep 22, 134–149 (2023).

11. Rosenberg, H., Pollock, N., Schiemann, A., Bulger, T. & Stowell, K. Malignant hyperthermia: a review. Orphanet J Rare Dis 10, 93 (2015).

12. Marty, I. & Fauré, J. Excitation-Contraction Coupling Alterations in Myopathies. J Neuromuscul Dis 3, 443–453 (2016).

13. Hopkins, P. M. et al. European Malignant Hyperthermia Group guidelines for investigation of malignant hyperthermia susceptibility. Br J Anaesth 115, 531–539 (2015).

14. Roux-Buisson, N. et al. Identification of variants of the ryanodine receptor type 1 in patients with exertional heat stroke and positive response to the malignant hyperthermia in vitro contracture test. Br J Anaesth 116, 566–568 (2016).

15. Bosson, C. et al. Variations in the TRPV1 gene are associated to exertional heat stroke. J Sci Med Sport 23, 1021–1027 (2020).

16. Endo, Y. et al. Variants in ASPH cause exertional heat illness and are associated with malignant hyperthermia susceptibility. Nat Commun 13, 3403 (2022).

17. Hopkins, P. M., Ellis, F. R. & Halsall, P. J. Evidence for related myopathies in exertional heat stroke and malignant hyperthermia. Lancet 338, 1491–1492 (1991).

18. Köchling, A., Wappler, F., Winkler, G. & Schulte am Esch, J. S. Rhabdomyolysis following severe physical exercise in a patient with predisposition to malignant hyperthermia. Anaesth Intensive Care 26, 315–318 (1998).

19. Dolphin, A. C. Voltage-gated calcium channel α 2δ subunits: an assessment of proposed novel roles. F1000Res 7, F1000 Faculty Rev-1830 (2018).

20. Wu, J. et al. Structure of the voltage-gated calcium channel Cav1.1 complex. Science 350, aad2395 (2015).

21. Lee, A. et al. Characterization of Cav1.4 complexes (α11.4, β2, and α2δ4) in HEK293T cells and in the retina. J Biol Chem 290, 1505–1521 (2015).

22. Dolphin, A. C. Biochemistry and physiology of voltage-gated calcium channel trafficking: a target for gabapentinoid drugs. Open Biol 15, 250013 (2025).

23. Qin, N., Yagel, S., Momplaisir, M.-L., Codd, E. E. & D’Andrea, M. R. Molecular cloning and characterization of the human voltage-gated calcium channel alpha(2)delta-4 subunit. Mol Pharmacol 62, 485–496 (2002).

24. Klomp, A. et al. The voltage-gated Ca2+ channel subunit α2δ-4 regulates locomotor behavior and sensorimotor gating in mice. PLoS One 17, e0263197 (2022).

25. Kerov, V. et al. α2δ-4 Is Required for the Molecular and Structural Organization of Rod and Cone Photoreceptor Synapses. J Neurosci 38, 6145–6160 (2018).

26. Wang, Y. et al. The Auxiliary Calcium Channel Subunit α2δ4 Is Required for Axonal Elaboration, Synaptic Transmission, and Wiring of Rod Photoreceptors. Neuron 93, 1359–1374.e6 (2017).

27. Cantí, C. et al. The metal-ion-dependent adhesion site in the Von Willebrand factor-A domain of alpha2delta subunits is key to trafficking voltage-gated Ca2+ channels. Proc Natl Acad Sci U S A 102, 11230–11235 (2005).

28. Briot, J. et al. A three-way inter-molecular network accounts for the CaVα2δ1-induced functional modulation of the pore-forming CaV1.2 subunit. J Biol Chem 293, 7176–7188 (2018).

29. Polster, A., Perni, S., Bichraoui, H. & Beam, K. G. Stac adaptor proteins regulate trafficking and function of muscle and neuronal L-type Ca2+ channels. Proc Natl Acad Sci U S A 112, 602–606 (2015).

30. Angelotti, T. & Hofmann, F. Tissue-specific expression of splice variants of the mouse voltage-gated calcium channel alpha2/delta subunit. FEBS Lett 397, 331–337 (1996).

31. Felix, R., Gurnett, C. A., De Waard, M. & Campbell, K. P. Dissection of functional domains of the voltage-dependent Ca2+ channel alpha2delta subunit. J Neurosci 17, 6884–6891 (1997).

32. Lefebvre, R., Pouvreau, S., Collet, C., Allard, B. & Jacquemond, V. Whole-cell voltage clamp on skeletal muscle fibers with the silicone-clamp technique. Methods Mol Biol 1183, 159–170 (2014).

33. Kutchukian, C. et al. Phosphatidylinositol 3-kinase inhibition restores Ca2+ release defects and prolongs survival in myotubularin-deficient mice. Proc Natl Acad Sci U S A 113, 14432–14437 (2016).

34. Obermair, G. J. et al. The Ca2+ channel alpha2delta-1 subunit determines Ca2+ current kinetics in skeletal muscle but not targeting of alpha1S or excitation-contraction coupling. J Biol Chem 280, 2229–2237 (2005).

35. Gong, H. C., Hang, J., Kohler, W., Li, L. & Su, T. Z. Tissue-specific expression and gabapentin-binding properties of calcium channel alpha2delta subunit subtypes. J Membr Biol 184, 35–43 (2001).

36. Murray, K. O. et al. Exertional heat stroke causes long-term skeletal muscle epigenetic reprogramming, altered gene expression, and impaired satellite cell function in mice. Am J Physiol Regul Integr Comp Physiol 326, R160–R175 (2024).

37. Ba-Abbad, R. et al. Mutations in CACNA2D4 Cause Distinctive Retinal Dysfunction in Humans. Ophthalmology 123, 668–671.e2 (2016).

38. Sambuughin, N. et al. Genetics of Exertional Heat Illness: Revealing New Associations and Expanding Heterogeneity. Int J Mol Sci 25, 11269 (2024).

39. Whittaker, C. A. & Hynes, R. O. Distribution and evolution of von Willebrand/integrin A domains: widely dispersed domains with roles in cell adhesion and elsewhere. Mol Biol Cell 13, 3369–3387 (2002).

40. Dahimene, S. et al. Biallelic CACNA2D1 loss-of-function variants cause early-onset developmental epileptic encephalopathy. Brain 145, 2721–2729 (2022).

41. Chen, Z. et al. EMC chaperone-CaV structure reveals an ion channel assembly intermediate. Nature 619, 410–419 (2023).

42. Dolphin, A. C. & Lee, A. Presynaptic calcium channels: specialized control of synaptic neurotransmitter release. Nat Rev Neurosci 21, 213–229 (2020).

43. Hoppa, M. B., Lana, B., Margas, W., Dolphin, A. C. & Ryan, T. A. α2δ expression sets presynaptic calcium channel abundance and release probability. Nature 486, 122–125 (2012).

44. Krawczak, M., Reiss, J. & Cooper, D. N. The mutational spectrum of single base-pair substitutions in mRNA splice junctions of human genes: causes and consequences. Hum Genet 90, 41–54 (1992).

45. Costa, J. G. et al. Sex differences in the thermoregulatory and cardiovascular response to exercise in hot environmental conditions. Am J Physiol Regul Integr Comp Physiol 329, R651–R660 (2025).

46. Alele, F., Malau-Aduli, B., Malau-Aduli, A. & Crowe, M. Systematic review of gender differences in the epidemiology and risk factors of exertional heat illness and heat tolerance in the armed forces. BMJ Open 10, e031825 (2020).

47. Cross-Disorder Group of the Psychiatric Genomics Consortium. Identification of risk loci with shared effects on five major psychiatric disorders: a genome-wide analysis. Lancet 381, 1371–1379 (2013).

48. van Loo, K. M. J. et al. Calcium Channel Subunit α2δ4 Is Regulated by Early Growth Response 1 and Facilitates Epileptogenesis. J Neurosci 39, 3175–3187 (2019).

49. Teixeira, M. et al. Electroporation of mice zygotes with dual guide RNA/Cas9 complexes for simple and efficient cloning-free genome editing. Sci Rep 8, 474 (2018).

