## Supplementary material for "CACNA2D4 variants are associated with exertional heat stroke susceptibility": supp figures

### Supplemental Figures

**Sup Fig.1 Membrane localization of human α2δ4 by immunofluoresence**
COS7 cells transfected with plasmids encoding human HA-α2δ4 (FL) or HA-α2δ4 S299R (FL S299R), labelled with anti-HA, anti-Cdk2 antibodies and AlexaFLuo488 WGA.
A/ Antibody labelling was performed after PFA fixation without permeabilization. Scale bar = 10 μm
B/ Antibody labelling was performed with PFA+ Saponin permeabilization. Scale bar = 20 μm

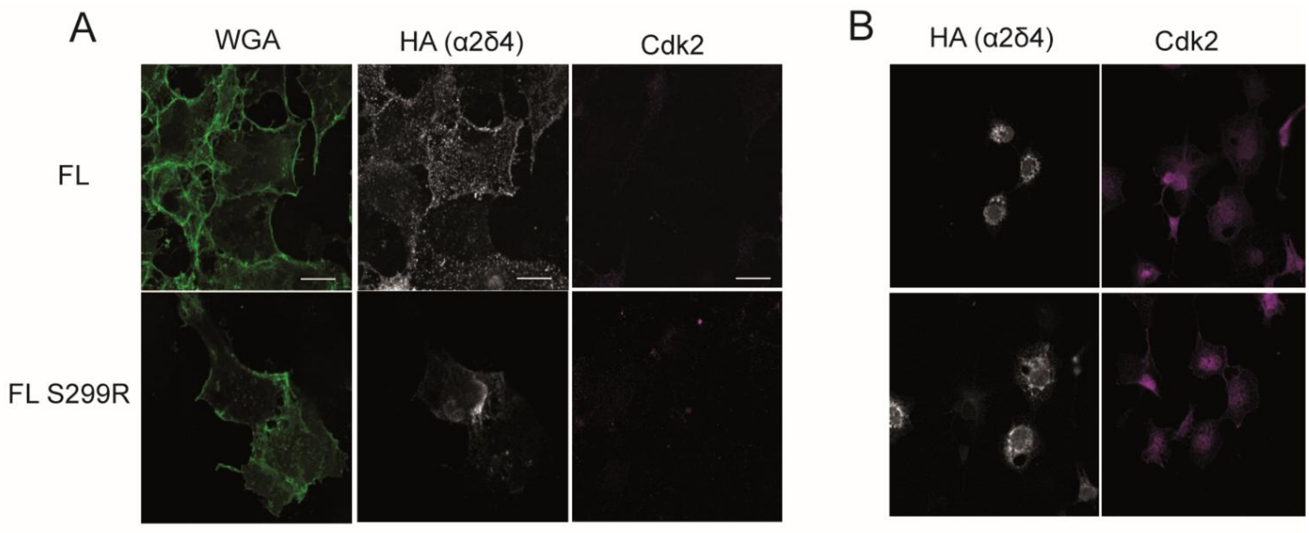


**Sup Fig.2 Co-immunoprecipitation of α1s with skm*CACNA2D4***
HEK293 cells were transfected with plasmids encoding tGFP- α1s + Flag-β1a subunits alone (α1s+β1a) or together with mouse full length (FL) or skeletal forms (skm) of HA-2δ4. After immunoprecipitation with anti HA antibody, lysate and immunoprecipitated fraction were recovered and probed with anti-GFP (α1s), anti-HA antibody (α2δ4) and anti-GAPDH antibody. NT= cells not transfected.

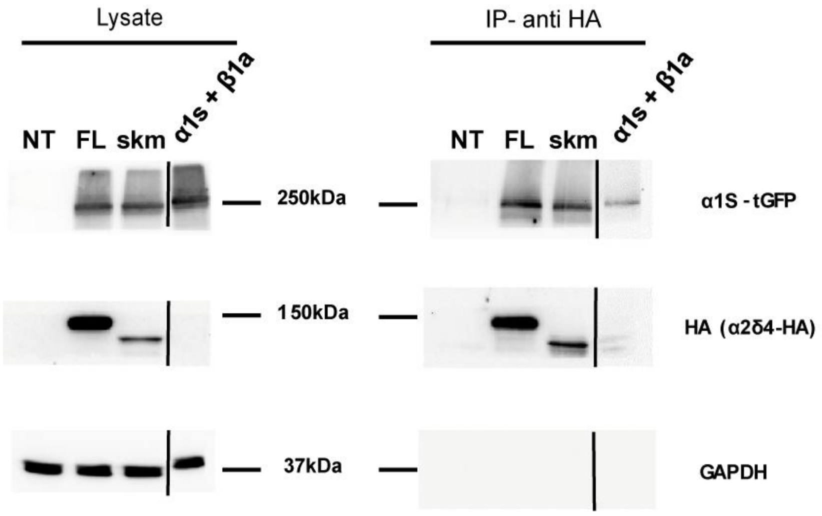

**Sup Fig.3 localization of mouse α2δ4**

COS7 cells transfected with plasmids encoding mouse HA-tagged full length α2δ4 (FL), HA-tagged skm-α2δ4 (skm), or HA tagged skm-α2δ4ΔCt (skmΔCT). Cells were labelled with anti HA or anti Climp63 antibodies Scale bar = 10 μm

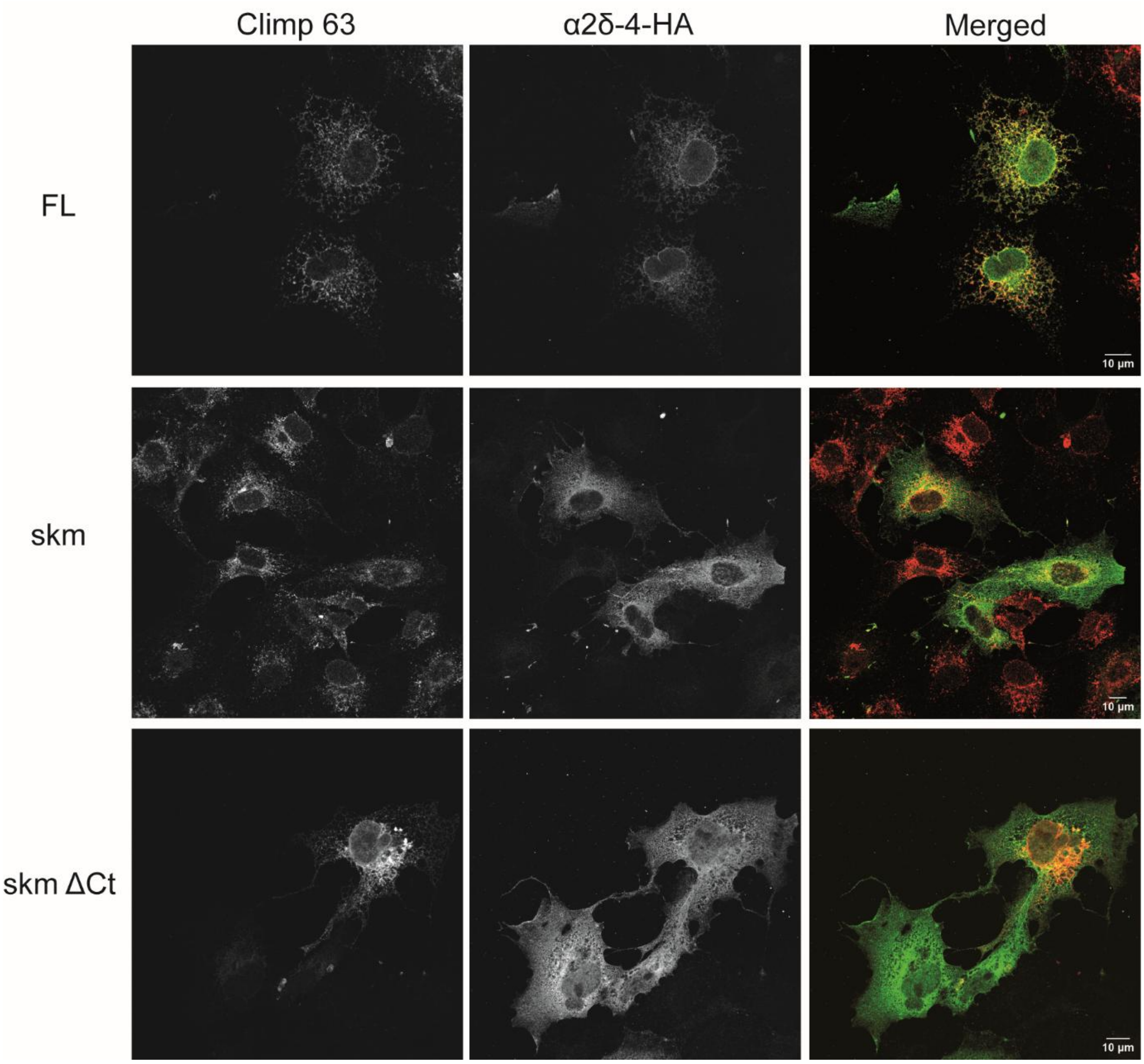

**Sup Fig.4 Expression of α2δ4 in WT and KO-*CACNA2D4* eye extracts.** Proteins were extracted from eye of WT or KO-*CACNA2D4* mice and separated by electrophoresis. Western-blot was performed with anti-α2δ4 and anti GAPDH antibodies. Arrows show 2 bands at the molecular weight of α2δ4 absent in extract from the *CACNA2D4* KO extract.

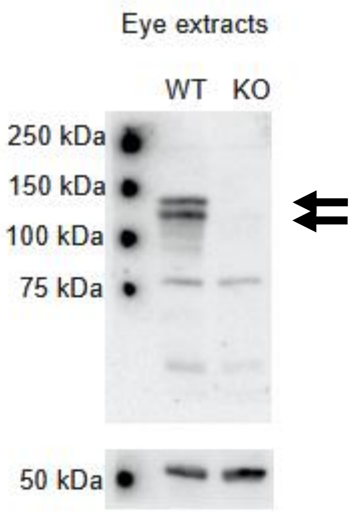
